# Sofosbuvir protects Zika virus-infected mice from mortality, preventing short- and long-term sequela

**DOI:** 10.1101/129197

**Authors:** André C. Ferreira, Camila Z. Valle, Patricia A. Reis, Giselle Barbosa-Lima, Yasmine Rangel Vieira, Mayara Mattos, Priscila de Paiva Silva, Carolina Sacramento, Hugo C. de Castro Faria Neto, Loraine Campanati, Amilcar Tanuri, Karin Brüning, Fernando A. Bozza, Patrícia T. Bozza, Thiago Moreno L. Souza

## Abstract

Zika virus (ZIKV) caused significant public health concern, because of its association with congenital malformations, neurological disorders in adults and, more recently, with deaths. Considering the necessity to mitigate the cases ZIKV-associated diseases, antiviral interventions against this virus are an urgent necessity. Sofosbuvir, a drug in clinical use against Hepatitis C Virus (HCV), is among the FDA-approved substances endowed with anti-ZIKV activity. In this work, we further investigated the *in vivo* activity of sofosbuvir against ZIKV. Neonatal Swiss mice were infected with ZIKV (2 x 10^7^ PFU) and treated with sofosbuvir at 20 mg/kg/day, a concentration compatible with pre-clinical development of this drug. We found that sofosbuvir reduced acute levels of ZIKV from 60 to 90 % in different anatomical compartments, such as in blood plasma, spleen, kidney and brain. Early treatment with sofosbuvir doubled the percentage and time of survival of ZIKV-infected animals, despite the aggressive virus challenge assayed and also prevented the acute neuromotor impairment triggered by the virus. On the long-term behavior analysis of ZIKV-associated sequelae, sofosbuvir prevented loss of hippocampal- and amygdala-dependent memory. Our results point out that sofosbuvir inhibits ZIKV replication *in vivo*, which is consistent with the prospective necessity of antiviral drugs to treat ZIKV-infected individuals.

## INTRODUCTION

Zika virus (ZIKV) is an enveloped,positive-sense single stranded RNA pathogen, which belongs to the *Flaviviridae* family. ZIKV is transmitted by mosquitoes, as other arboviruses of the *Flavivirus* genus. ZIKV re-emerged in the last few years and revealed to be a unique pathogen^1^. It spread through sexual and physical contact, and also vertically^1^. The main wave of Zika epidemics in Americas occurred from the middle of 2015 to the beginning of 2016, when the World Health Organization (WHO) declared that this outbreak was a public health emergency of international concern. This relevant apprehension was raised due to ZIKV association with congenital malformations, including microcephaly, and a broad range of neurological disorders in adults, including Guillain-Barré syndrome^2,3^. With the number of cases rising, Zika-associated deaths have also been reported^4,5^. Considering the necessity to mitigate the cases of ZIKV-associated morbidities, antiviral interventions against ZIKV are an urgent necessity.

Different studies have been published repurposing Food and Drug Administration (FDA)-approved molecules to treat ZIKV infection^6-11^. Over 30 FDA-approved molecules are endowed with anti-ZIKV activity. Among them, sofosbuvir, a clinically approved anti-hepatitis C virus (HCV), directly inhibits ZIKV RNA polymerase^12-14^. Like in HCV, we and others demonstrated that sofosbuvir targets ZIKV RNA polymerase, leading to inhibition of virus replication in cellular types important for the pathogenesis of this emergent agent, such as human brain organoids, neural stem cells and neuroepithelial stem cells^12-14^. Therefore, sofosbuvir may represent a very selective option to treat Zika. Nevertheless, more detailed analyses of sofosbuvir’s anti-ZIKV activity *in vivo* are necessary.

Different animal models of Zika virus infection were reported recently^15^. It is considered that many of these models, from immunocompromised to immunocompetent neonatal mice, among others, are relevant for antiviral testing^15^. In general, for pharmacological studies, outbred animals, such as Swiss mice, represent a consistent model. These animals display broader responses, often found in heterogeneous population^16^, such as humans. Indeed, ZIKV infection in Swiss neonatal mouse models has been characterized, since 1950 ^15,17-19^. Another advantage of using neonatal mouse models is the opportunity to further examine behavioral sequelae provoked by the infection and the ability of testing specific treatments to overcome/prevent this phenotype.

In this study, we show that sofosbuvir protects ZIKV-infected animals from mortality by a very aggressive virus challenge. This was associated with a decrease in the viral RNA levels in different tissues as well as prevented of sequelae to the neuromotor system and to animal memory.

## MATERIAL AND METHODS

### Reagents

The antiviral drug sofosbuvir (β-d-2’-deoxy-2’-α-fluoro-2’-β-C-methyluridine) was donated by the BMK Consortium: Blanver Farmoquímica Ltda; Microbiológica Química e Farmacêutica Ltda; Karin Bruning & Cia. Ltda, (Taboão da Serra, São Paulo, Brazil). Drugs were dissolved in 100 % dimethylsulfoxide (DMSO) 1:10 (mass/volume), followed by appropriated dilutions in PBS or culture medium (DMEM) to treat the animals. The final DMSO concentrations showed no toxicity to the animals. Other materials were purchased from Thermo Scientific Life Sciences (Grand Island, NY), unless otherwise mentioned.

### Cells

African green monkey kidney (Vero) cells were cultured in DMEM. The culture medium was supplemented with 10 % fetal bovine serum (FBS; HyClone, Logan, Utah), 100 U/mL penicillin, and 100 μg/mL streptomycin. Cells were kept at 37 °C in 5 % CO_2_.

### Virus

ZIKV (MR766 strain) was propagated and titered by plaque-forming assays in Vero cells. Passages were carried out at a multiplicity of infection (MOI) of 0.01 at 26 °C. for 1 h at 37 °C. Next, the residual virus particles were removed by washing with phosphate-buffered saline (PBS), and the cells were cultured for additional 5 days. After each period, the cells were lysed by freezing and thawing, and centrifuged at 1,500 x *g* at 4 °C for 20 min to remove cellular debris. The virus was stored at -70 °C for further studies.

### Plaque-forming assay

Monolayers of Vero cells in 6-well plates were exposed to different dilutions of the supernatant containing virus for 1 h at 37 °C. Next, the cells were washed with PBS, and DMEM containing 1 % FBS and 3 % carboxymethylcellulose (Fluka) (overlay medium) was added to cells. After 5 days at 37 °C, the monolayers were fixed with 10 % formaldehyde in PBS and stained with a 0.1 % solution of crystal violet in 70 % methanol, and the virus titers were calculated by scoring the plaque-forming units (PFU).

### Molecular detection of virus RNA levels

Total RNA from culture, extract containing organs in PBS or plasma was extracted using QIAamp Viral RNA or RNeasy Mini Kits (Qiagen®), according to manufacturer’s instructions. Quantitative RT-PCR was performed using QuantiTect or QuantiNova Probe RT-PCR Kit (Quiagen®) in an ABI PRISM 7300 Sequence Detection System (Applied Biosystems). Amplifications were carried out in 25 μL reaction mixtures containing 2× reaction mix buffer, 50 μM of each primer, 10 μM of probe and 5 μL of RNA template. Primers, probes and cycling conditions recommended by the Centers for Disease Control and Prevention (CDC) protocol were used to detect the ZIKV^20^. The standard curve method was employed for virus quantification. For reference on the cell amounts used, the housekeeping gene RNAse P was amplified^20^. The Ct values for this target were compared to those obtained to different cell amounts, 10^7^ to 10^2^, for calibration.

### Animals

Swiss albino mice (*Mus musculus*) (pathogen free) from the Oswaldo Cruz Foundation breeding unit (Instituto de Ciência e Tecnologia em Biomodelos (ICTB)/Fiocruz) were used for these studies. The animals were kept at a constant temperature (25 °C) with free access to chow and water and a 12-h light/dark cycle. The Animal Welfare Committee of the Oswaldo Cruz Foundation (CEUA/FIOCRUZ) approved and covered (license number L-016/2016) the experiments in this study. The procedures described in this study were in accordance with the local guidelines and guidelines published in the National Institutes of Health Guide for the Care and Use of Laboratory Animals. The study is reported in accordance with the ARRIVE guidelines for reporting experiments involving animals^21^. The experimental laboratory received pregnant mice (at approximately the 14th gestational day) from the breeding unit. Pregnant mice were observed daily until delivery, to accurately determine the postnatal day. We established a litter size of 10 animals for all experimental replicates.

### Experimental infection and treatment

3-days-old Swiss mice were infected intraperitoneally with 2 x 10^7^ PFU of virus^22^. Treatments with sofosbuvir were carried out with sofosbuvir at 20 mg/kg/day intraperitoneally. Treatment started one day prior to infection (pre-treatment) or two days after infection (late treatment). Either way, treatment was conducted for 7 days. For comparisons, mock-infected and -treated groups of animals were used as controls. Animals were monitored daily for survival, weight gain and virus-induced short-term sequelae (righting in up to 60 seconds) (Supplementary Video). Animals were euthanized, when differences in weight gain between infected and control groups were > 50% and/or severe illness present. Blood was collected by cardiac puncture and placed on citrate-containing tubes for plasma separation. Tissues (spleen, kidney and brain) were collected. Initially, the tissues were analysed macroscopically, for the presence of pathological signs. Whenever possible, the pathological signs were quantified by counting per verified tissue/organs. Alternatively, tissues were lysed (RLT buffer; Qiagen) and homogenized with Potter-Elvehjem homogenizer (Teflon pestle and glass mortar). Homogenates were cleared by centrifugation and total RNA was extracted.

### Behavioral tests

To test the righting reflex, animals were tested daily during the course of acute infection. Animals were held in a supine position with all four paws facing up in the air for 5 seconds. After that, animals were released and the time the animal took to flip over onto its stomach with all four paws touching the surface was measured. A maximum of 60 seconds was given for each trial and animals were tested twice a day with a 5 minutes minimum interval between trials. For each animal, the lowest time was plotted in the graph. Animals that failed the test were included in the graph with a time of 60 seconds. Please see the Supplementary Video.

Animals at age of 6 to 8 weeks of old were assayed. The Morris water maze (MWM) is a behavioral task to evaluate hippocampal dependent learning and spatial memory. The water maze comprised a black circular pool (100 cm in diameter) that was conceptually divided into four equal, but imaginary, quadrants for the purpose of data analysis. The water temperature was 25 °C. A platform (10 cm^2^), which was hidden from the mouse’s view, was located 2 cm beneath the surface of the water, allowing the mouse to easily climb onto it once its presence was detected. The water maze was located in a well-lit white room with several posters and other distal visual stimuli hanging on the walls to provide spatial cues. Training on the hidden platform (spatial) version of the MWM was carried out on 4 consecutive days. On day 5, when the platform is removed, memory is evaluated.

An amygdala-dependent aversive memory assay (freezing test) was conducted in a chamber with 3 dark walls and clear frontal wall and lid (28 x 26 x 23 cm). The floor of the chamber consisted of 20 parallel stainless steel grid bars, each measuring 4 mm in diameter and spaced 7 cm apart. The grid was connected to a shock generator device (Insight LTDA, Brazil). Training session consisted of placing mouse in the chamber and allowing a 3 min acclimation period. After, mice received two foot-shocks (0.6 mA, 3 s with one interval of 30 s), and then were returned to their home cages. 24 hours later, mice were exposed to the same environment without shock stimuli, during 3 min. Memory was assessed and expressed as the percentage of time that mice spent freezing (considered when it was crouching without movement and the head except that associated with breathing).

### Statistical analysis

Significance of survival curves was evaluated using the Log-rank (Mantel-Cox) tests. Behavioral tests were analyzed with ANOVA, followed by Tukey’s post hoc test. Odds ratio (OR) and 95 % confidence interval (CI) were calculated by Fisher’s exact test with Lancaster’s mid-P correction. *P* values of 0.05 or less were considered statistically significant.

## RESULTS

### Sofosbuvir enhances the survival of ZIKV-infected mice

We infected 3-days old Swiss mice intraperitoneally with ZIKV at 2 X 10^7^ PFU ^22^, which represents a very aggressive virus challenge likely to drive near 100 % mortality within days, under our experimental conditions. Treatments with sofosbuvir were carried out at a dose of 20 mg/kg/day, which is compatible with the pre-clinical studies conducted for further clinical approval of this drug^23^. Animals were treated for 7 days, beginning at 1 day before infection or 2 days after infection (Figure 1 and Table 1).

**Figure 1.**
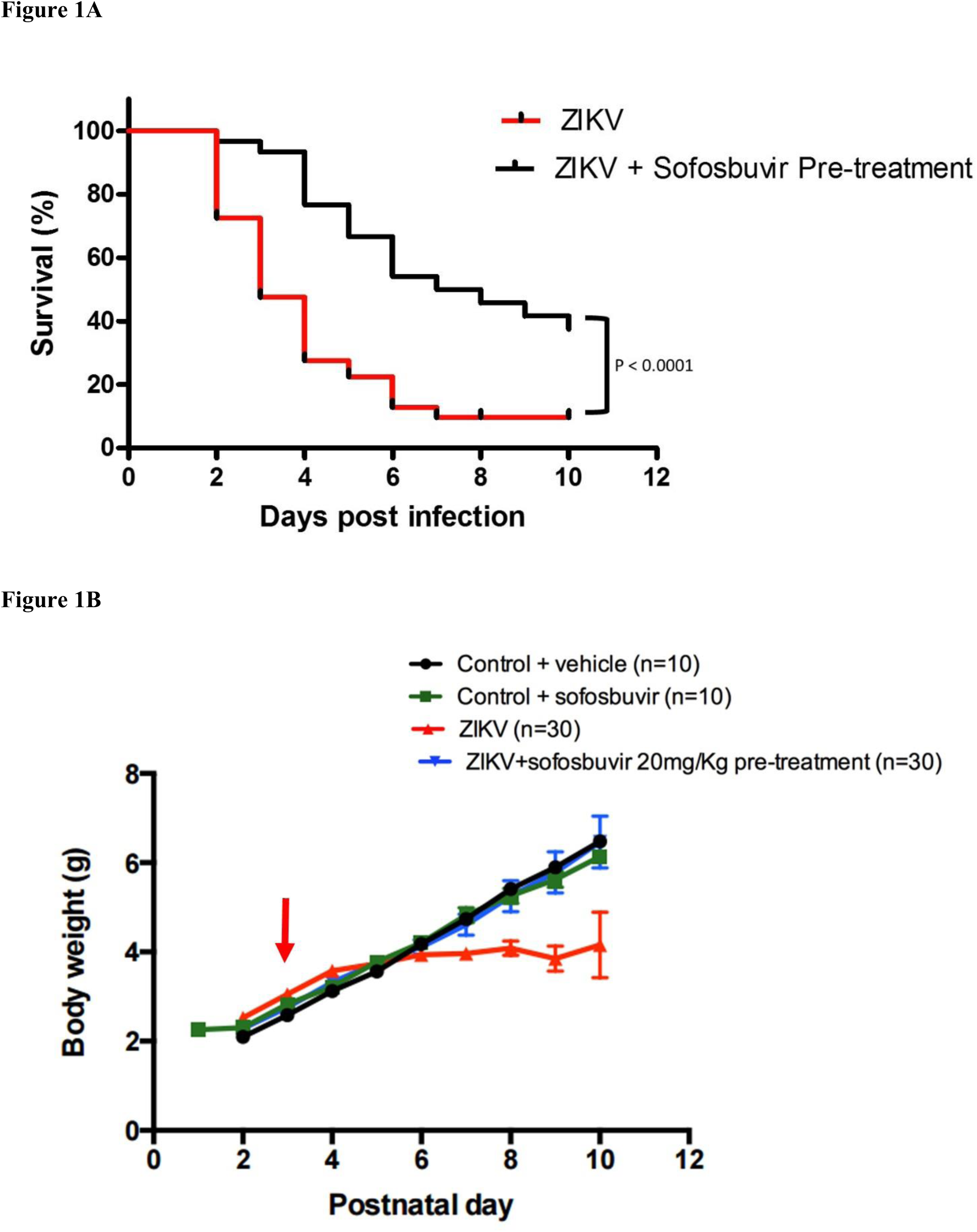
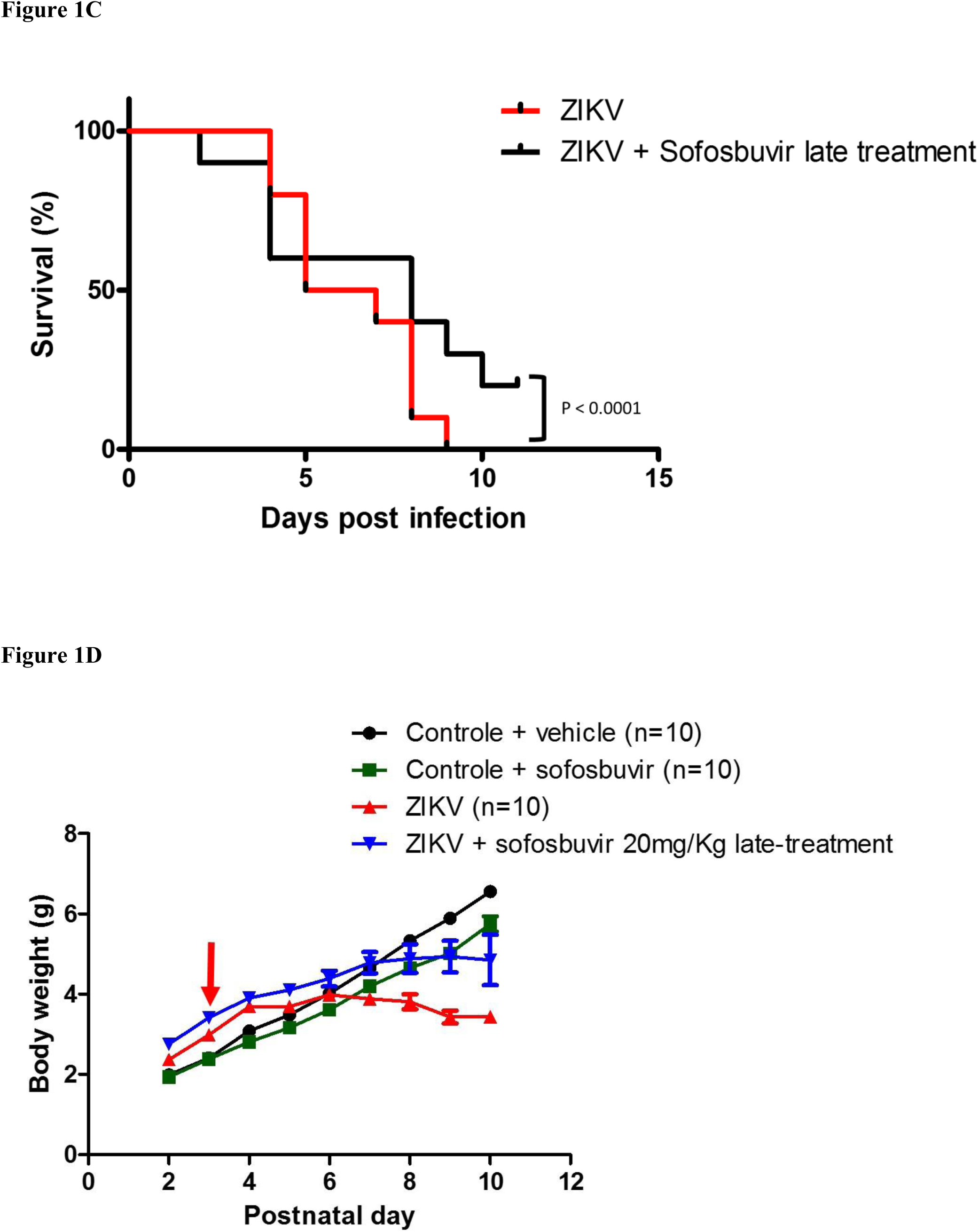
Treatment with Sofosbuvir increased survival and inhibited weight loss of ZIKV-infected mice. 3**-**days old Swiss mice were infected with ZIKV (2 x10^7^ PFU) and treated with sofosbuvir either 1 day before (A and B) or 2 days after infection (C and D). Survival (A and C) and weight variation (B and D) were assessed during the course of treatment (7 days). The red arrow indicates when animals were infected. Survival was statistically assessed by Log-rank (Mentel-Cox) test. Differences in weight is displayed as mean ± SEM, two-way ANOVA for each day was used to assess the significance. * P < 0.01; ** < 0.001.

**Table 1.**
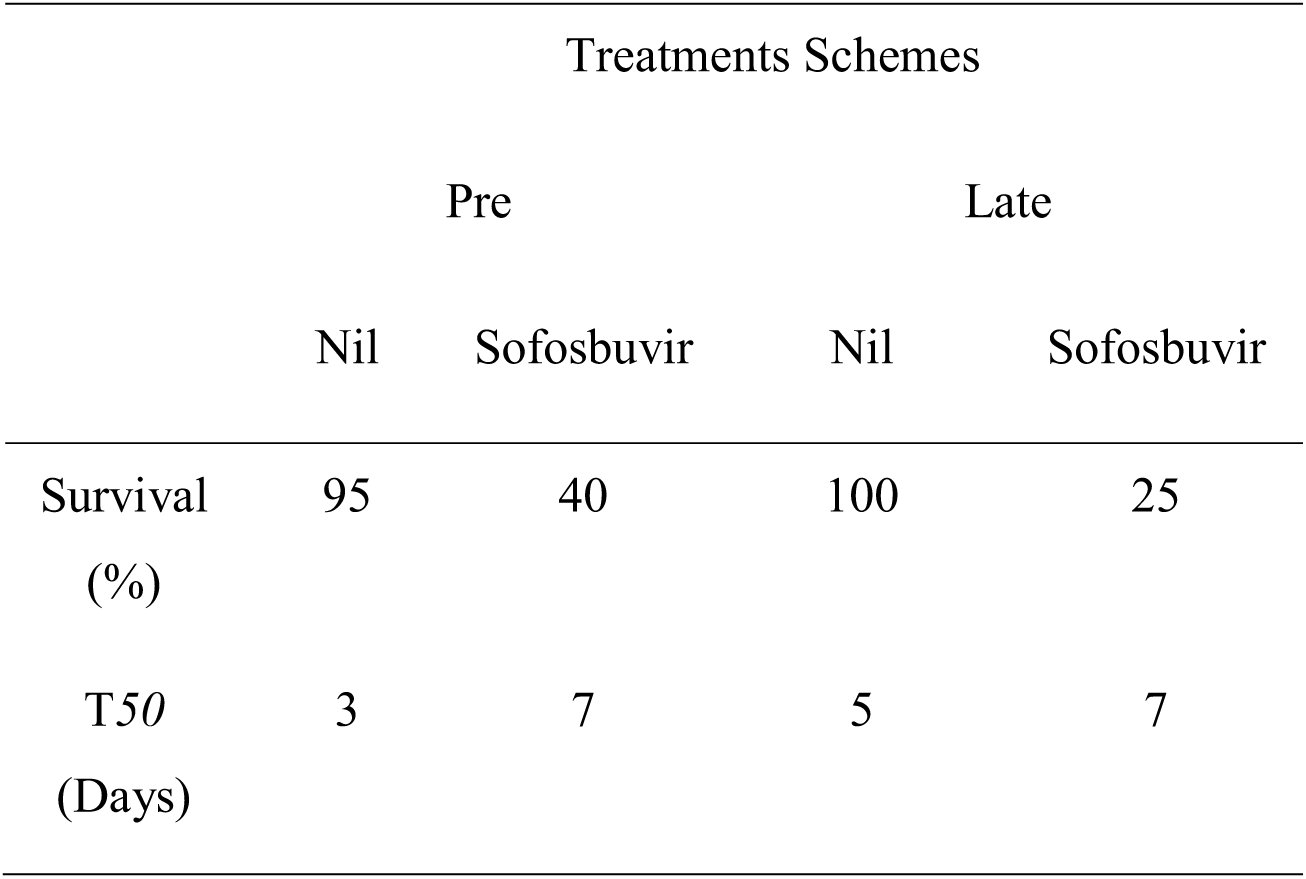
– Percentage and time of survival of sofosbuvir-treated ZIKV-infected mice

|  | Treatments Schemes |  |  |  |
| --- | --- | --- | --- | --- |
|  | Pre |  | Late |  |
|  | Nil | Sofosbuvir | Nil | Sofosbuvir |
| Survival<br>(%) | 95 | 40 | 100 | 25 |
| T50<br>(Days) | 3 | 7 | 5 | 7 |

Sofosbuvir significantly protected pre-treated infected mice from ZIKV-induced mortality (Fig 1A). Virtually all ZIKV-infected mice died up to 7 days after infection, whereas overall 40 % of sofosbuvir-treated ZIKV-infected animals survived (Fig 1A). Along with the survival curves, we evaluated the weight gain of pre-treated animals during the time course of the assay. ZIKV-infected animals have an impaired postnatal development, whereas sofosbuvir-treated ZIKV-infected mice weight gain was indistinguishable from uninfected controls (Fig 1B).

Postponing the treatment to the second day after infection still preserved some level of protection to ZIKV-infected mice (Fig 1C). All infected and untreated mice died at 8 days post infection, whereas 25 % of the mice receiving late treatment survived (Fig 1C). With respect to the postnatal development of ZIKV-infected mice, treated animals gained more weight than untreated mice (Fig 1D).

Remarkably, both treatment regimens resulted in enhanced survival rate when compared to the absence of treatment. Nevertheless, pretreatment may be considered more effective. Pretreatment doubled the increase in survival rates and the mean time of survival (T_*50*_), when compared to late treatments (Table 1). To statistically compare different experimental groups with respect to mortality risk, odds ratio (OR) for this outcome were calculated. Comparing no treatment with sofosbuvir treatment (irrespective of timing) represented a reduction in mortality risk (OR = 0.198) (Fig 2). Still comparing with no treatment, pre-treatment was more effective than late (no vs. pre: OR = 0.098; no vs. late OR = 0.220) (Fig 2). The comparison between late vs pre-treatment also revealed the benefits of the earlier interventions (OR = 0.648) (Fig 2).

**Figure 2.**
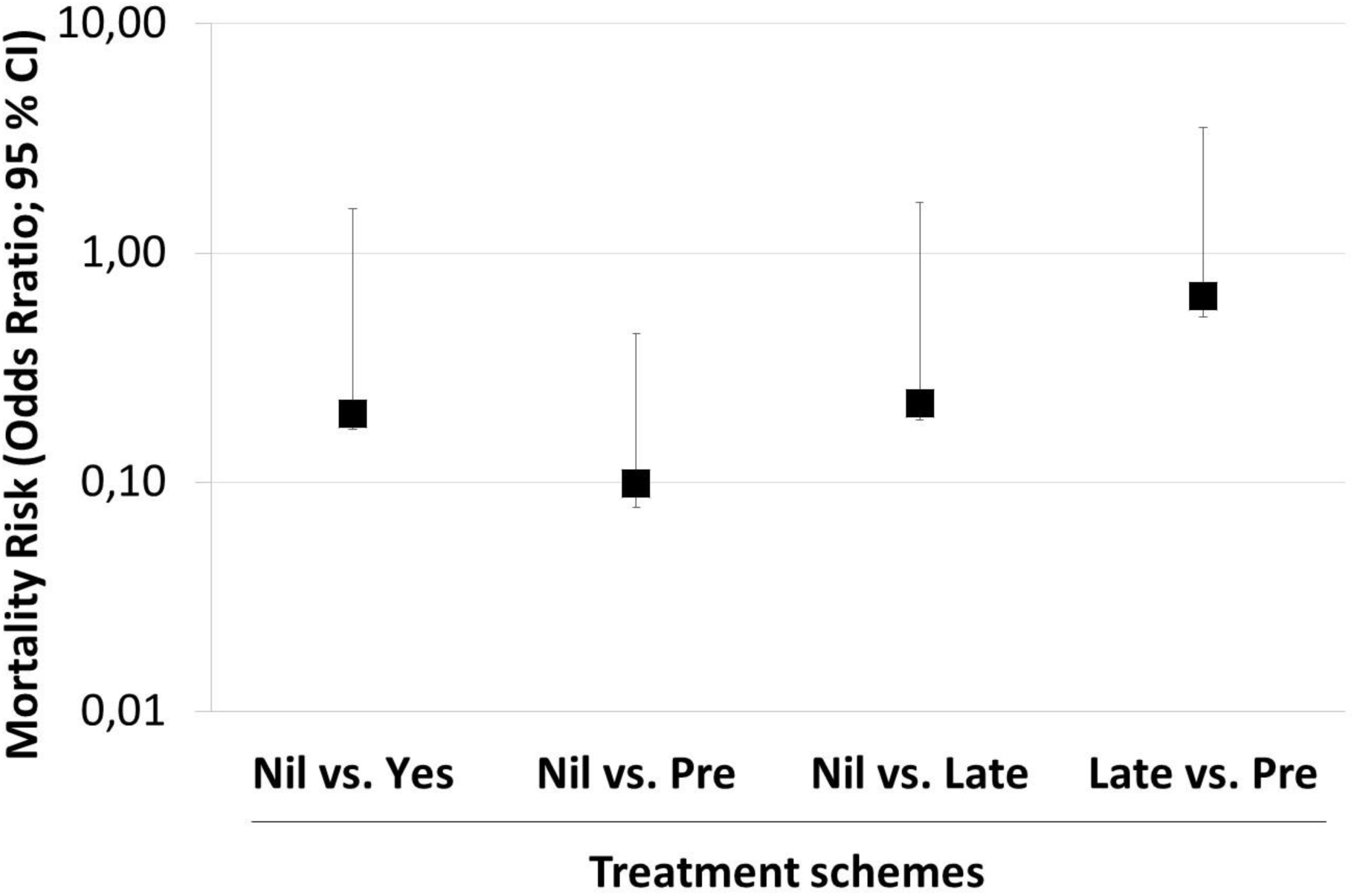
Sofosbuvir reduced the risk mortality of ZIKV-infected mice. Taking the data from Figures 1A and C as reference, odds ratio for mortality risk (with 95 % confidence interval; CI) was calculated by Fisher’s exact test with Lancaster’s mid-P correction. All comparisons were statistically significant, with P < 0.001.

These data may provide two important general notions: i) earlier sofosbuvir administration to ZIKV-infected mice leads to better antiviral results and to reduced mortality; and ii) regardless of timing, it is more opportune to administer sofosbuvir rather than leave the animals without treatment, for the sake of survival.

### Sofosbuvir decreases ZIKV loads during acute infection

Since sofosbuvir inhibits ZIKV replication, enhanced survival due to this treatment was presumably associated with reduction in viral levels during acute infection. We evaluated this hypothesis and measured the magnitude of virus inhibition *in vivo*. To do so, sofosbuvir-treated ZIKV-infected animals were euthanized daily from the first to the fifth day after infection. Next, viral loads were measured in different tissues (Fig 3). We observed that sofosbuvir reduced the mice viremia over 90 %, especially during the first 3 days after infection, when an exponential increase in viral levels in the plasma was observed (Fig 3A). Sofosbuvir reduced by up to 60 % the peak of virus detection between 2 and 4 days post-infection in the kidney (Fig 3B). Virus detection in the spleen and brain was abundant at 4 to 5 days after infection and sofosbuvir treatment reduced up to 80 % virus levels in these tissues (Fig 3C and D). Remarkably, during this experiment, we observed that ZIKV-infected animals had focci of cerebral microhemorrhage. Sofosbuvir prevented/reduced ZIKV-induced microhemorrhage over 90 % (Fig 4) in pretreated animals.

**Figure 3.**
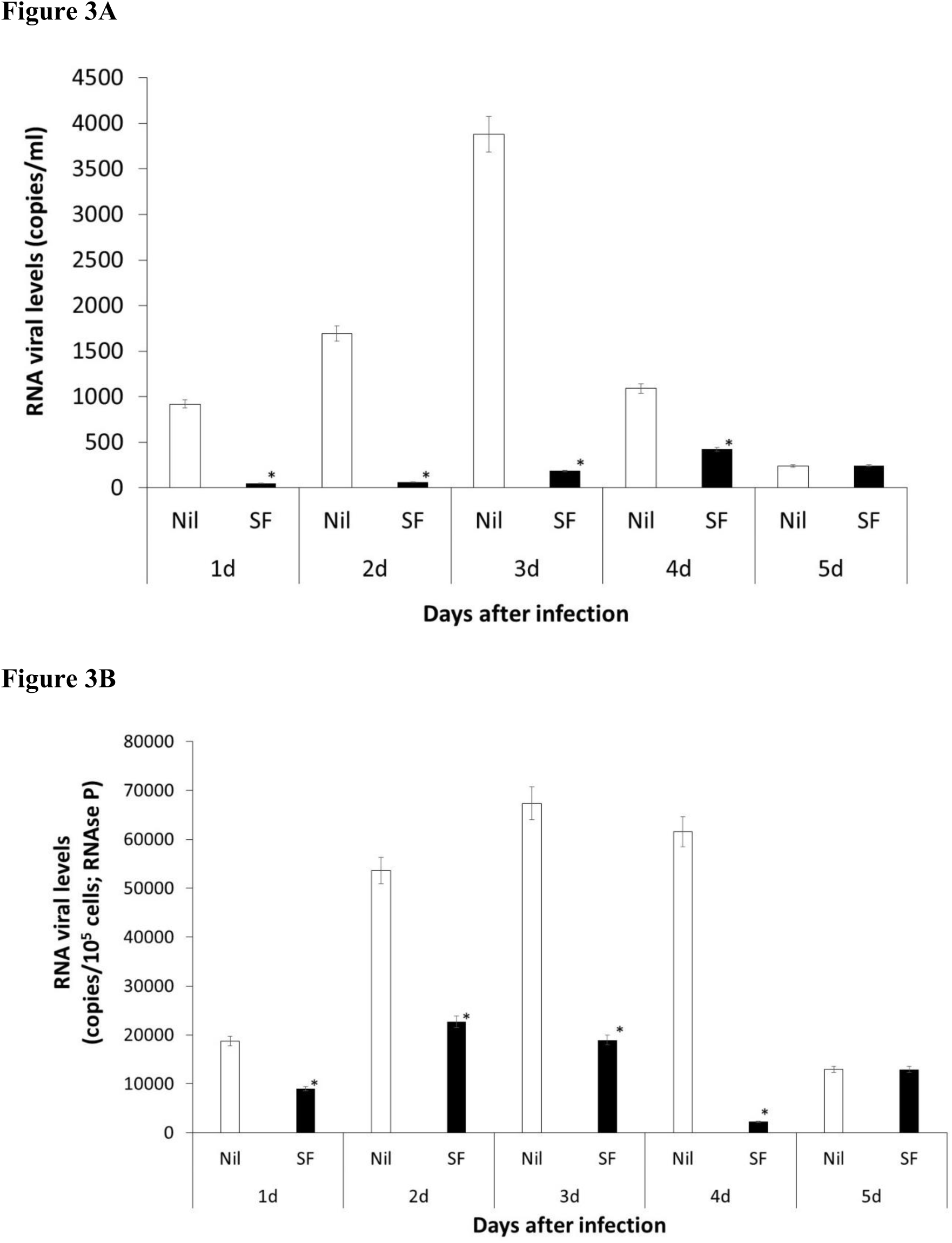
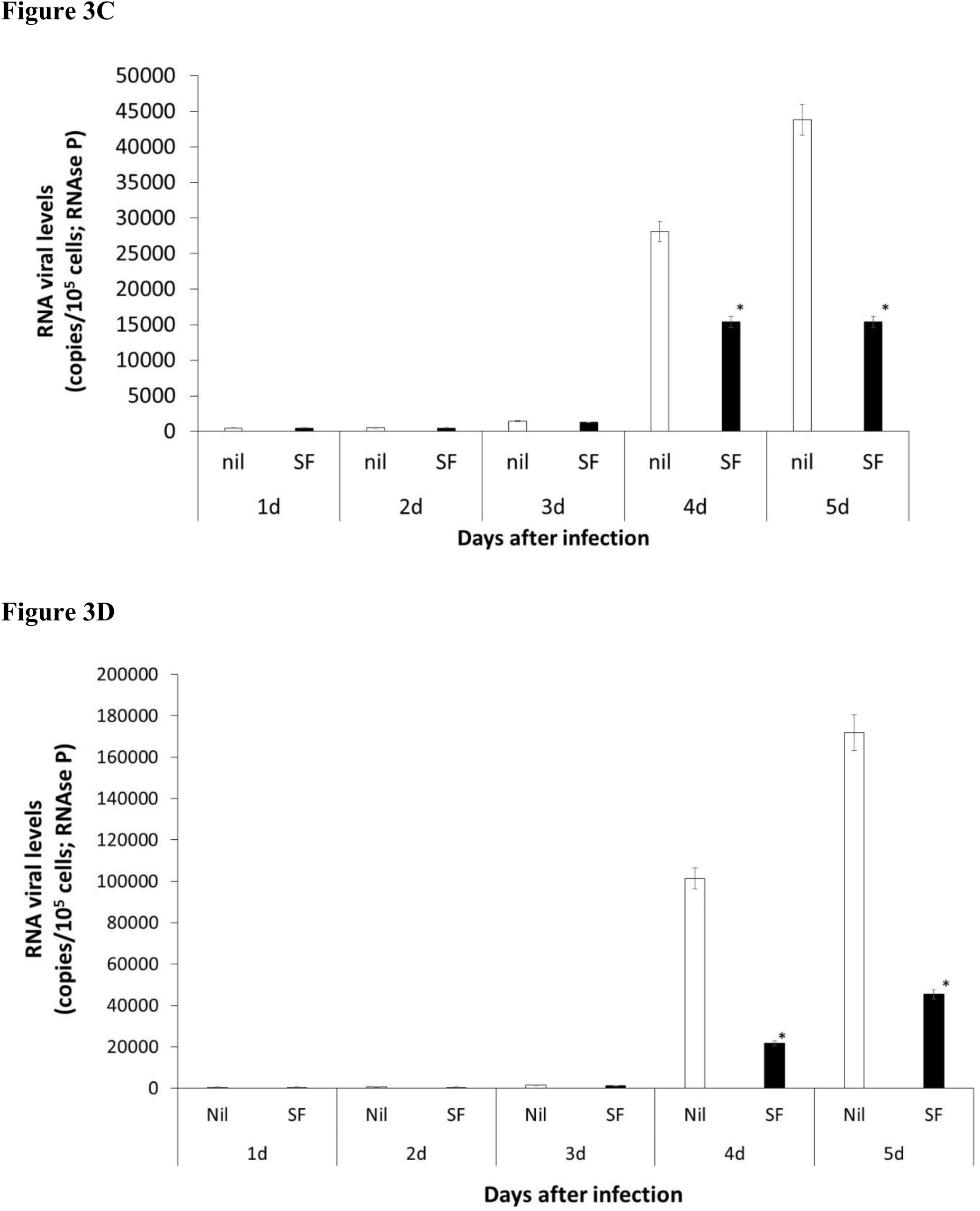
Sofosbuvir-dependent inhibition of ZIKV replication reduced viral loads in different anatomical compartments during acute infection. 3**-**days old Swiss mice were infected with ZIKV (2 x10^7^ PFU) and treated with with sofosbuvir (SF) or not (nil), starting at day 1 before the infection. At indicated days after infection, animals were euthanized and ZIKV RNA levels were measured in the plasma (A), kidney (B), spleen (C) and brain (D). Results are displayed as mean ± SEM. Student’s T test was used to compare the viral levels from SF- vs. mock-treated mice. * P < 0.01.

**Figure 4.**
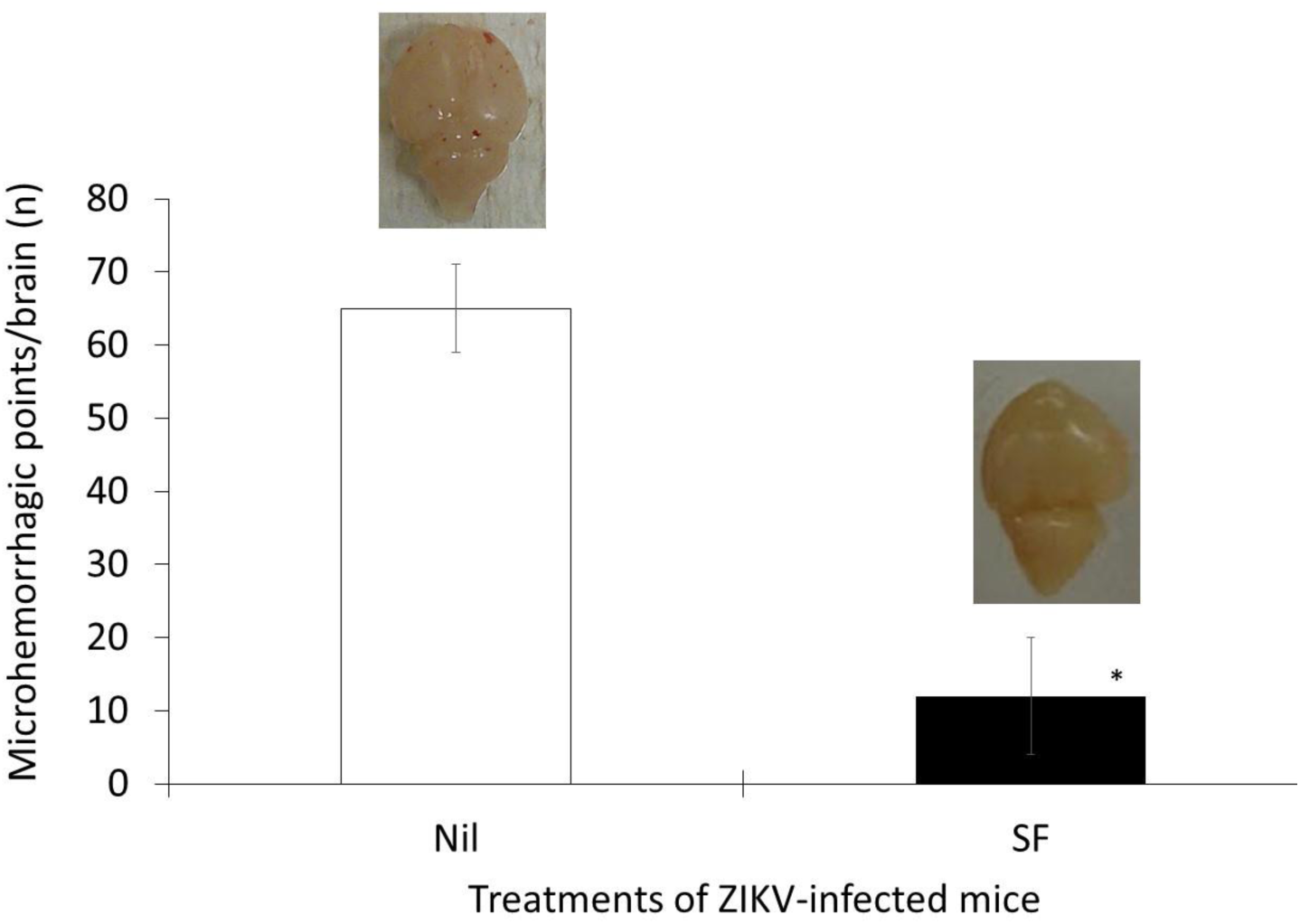
Sofosbuvir decreases the degree of microhemorrhage in the brain of ZIKV-infected mice. 3**-**days old Swiss mice were infected with ZIKV (2 x10^7^ PFU) and treated with sofosbuvir (SF) or not (nil) starting at day 1 before the infection. At the fifth day after infection animals were euthanized and whole brain collected to quantify the microhemorrhagic foci. Mean ± SEM is displayed. Student’s T test was used to compare the viral levels from SF- vs. mock-treated mice. * P < 0.01. The insets are representative brains of untreated and treated mice.

Altogether, our results indicate that sofosbuvir effects on mice survival was indeed followed by a reduction in virus detection in different anatomical compartments.

### Sofosbuvir prevents short- and long-term sequelae in ZIKV-infected mice

The neonatal animal model may represent a relevant model to evaluate short, and especially, long-term behavioral sequelae after infections. Consistently, we observed an acute neuromotor impairment in ZIKV-infected mice (Supplementary Video). To determine the magnitude of this injury and the benefits of sofosbuvir use, we applied the righting test reflex, for up to 60 seconds. ZIKV-infected animals, untreated with sofosbuvir, took over 12-fold more time to get to the upright position than sofosbuvir-treated animals or controls (Fig 5). No statistically significant differences were observed between sofosbuvir-treated ZIKV-infected mice and the controls (Fig 5). Our date indicate that sofosbuvir protected the animals from ZIKV-associated neuromotor impairment.

**Figure 5.**
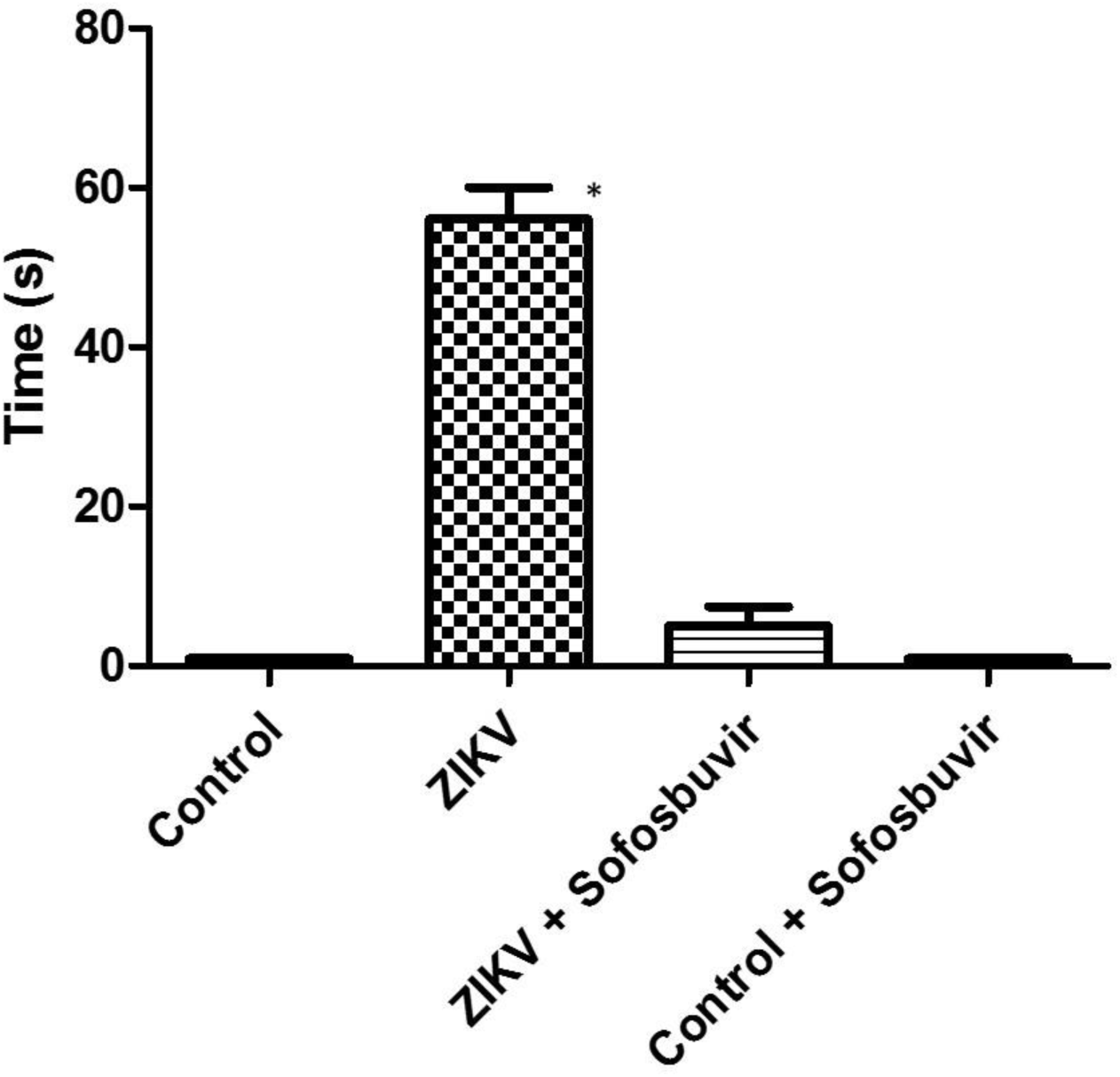
Sofosbuvir prevented neuromotor impairment in the ZIKV-infected mice. 3**-**days old Swiss mice were infected with ZIKV (2 x10^7^ PFU) and treated with sofosbuvir (SF) or not (nil) starting at day 1 before the infection. At the fifth day after infection animals were tested for righting (Supplementary Video). Animals were turned backwards and allowed up to 60 second to return to the upright position. Results are presented as mean ± SEM. Student’s T test was used to compare untreated ZIKV-infected mice with other groups individually. * P < 0.01.

Moreover, we had a few ZIKV-infected mice that did not succumb to the infection (Fig 1A). We kept these mice for 6 to 8-week to further monitor behavioral sequelae. We applied the Morris water maze test to assess hippocampal learning and memory. On the learning tests, training to find a platform 2 cm beneath the surface of the water was carried for 4 days. Healthy control animals and survivors from ZIKV infection responded similarly to learning, independently on whether infected animals were treated or not (Fig 6A). On day 5, the platform was removed and memory was evaluated, by measuring the timing to stay in the platform’s quadrant (latency). Untreated ZIKV-infected mice did not stay on the quadrant where the platform had been previously located, in comparison to control (uninfected) and sofosbuvir-treated ZIKV-infected mice (Fig 6B).

**Figure 6.**
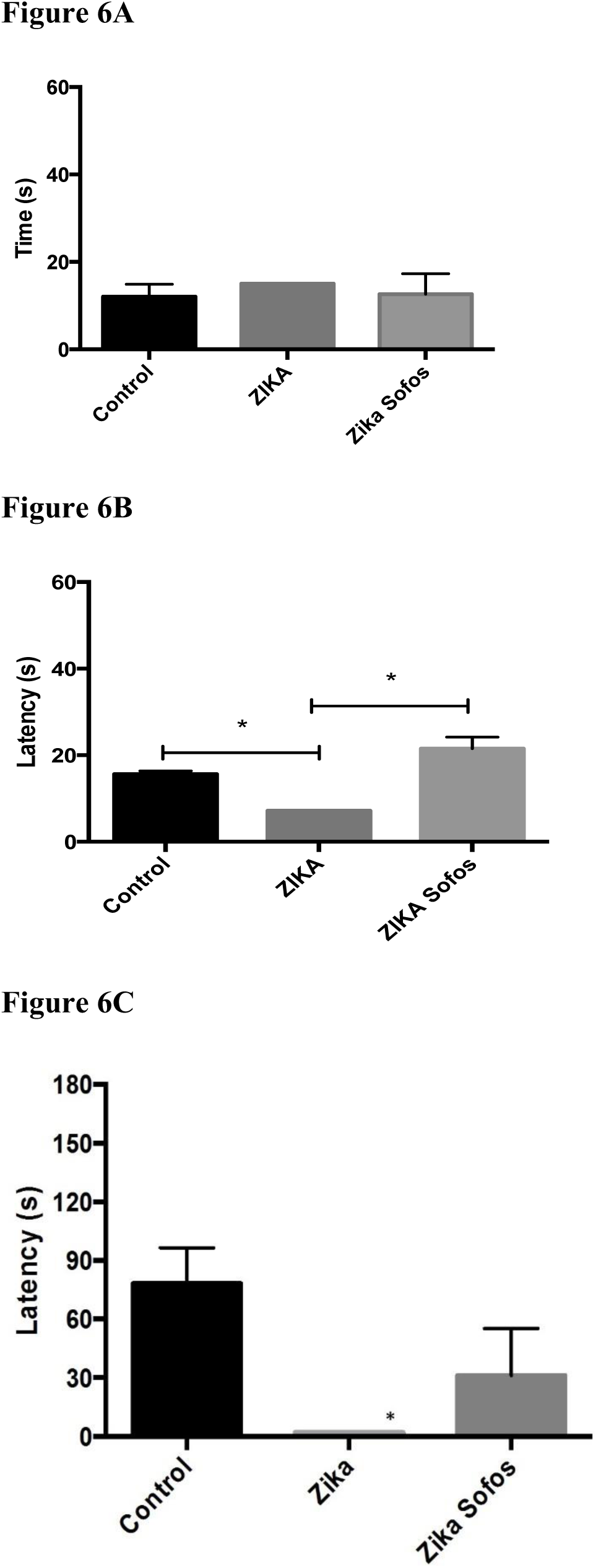
Sofosbuvir prevented memory loss in animals that survived from ZIKV infection. 3**-**days old Swiss mice were infected with ZIKV (2 x10^7^ PFU) and treated with sofosbuvir (SF) or not (nil) starting at day 1 before the infection. Animals that survived were kept and tested for behavior sequela in learning and memory after 60 days. Time to find the platform, according to MWM test, was performed (A). Trial in the absence of the platform was conducted (B) (*p<0.05, Student T test). In the panel B, latency represents the time spent exploring the quadrant where the platform was located before removal, and data are expressed as the mean ± SEM (n= 5-10). Aversive memory was evaluated by freezing behavior 24 h post-training session, whereas mice were allowed to explore the aversive environment during 180 followed by two foot-shock (0.6 mA, 3 s) (C). In panel C, latency represents the time spent without movement (freezing) during 180 s, and data are expressed as the mean ± SEM (n= 2-5).

Subsequently, an amygdala-dependent aversive memory test was performed (freezing test). This test consists of two foot-shocks on mice. On the next day, mice are exposed to the same environment, without shock, when latency is measured. Our data showed that untreated ZIKV-infected animal lost the aversive memory, whereas the sofosbuvir-treated ZIKV-infected mice and healthy controls behaved normally (Fig 6C). Our results indicate that besides the increase in survival lead by sofosbuvir, this drug also prevent from ZIKV-induced behavior sequelae, neuromotor impairments and memory loss.

## DISCUSSION

In the last years, the risk perception on ZIKV infection increased substantially. Although Zika fever is a mild and self-limited disease to most cases ^24^, ZIKV-associated morbidities have been described^2,3^. Since 2013 ZIKV spread explosively across immunologically naïve populations throughout the world, and especially in the Americas^25^. For instance, in Brazil during 2015, it is estimated that over 4 million people were affected by this virus^26^. Major concerns were raised due to the association of ZIKV infection with neurological disorders during fetal development and adulthood^2,3^. More recently, a few Zika-associated deaths have also been reported^4,5^. We and others have shown that sofosbuvir, a clinically approved drug against HCV, show strong antiviral activity against ZIKV^12-14^. Particularly, we showed that sofosbuvir is functionally active against ZIKV in cells derived from peripheral organs and the CNS, by targeting the viral RNA polymerase^13^. To advance on the pre-clinical development of sofosbuvir as an anti-ZIKV drug, we further examined whether this uridine analog is active *in vivo*.

We understand that pharmacological studies on animal models, such as Swiss outbred mice, with representative genetic heterogeneity, may allow further exploration of the generated data to a broader population^16^. For this reason, we have chosen to use these WT and outbred animal model for these analysis. Immunocompetent mice is a more natural model and may have limited susceptibility to ZIKV infection^15^, allowing some mice to survive even after an aggressive virus challenge. Furthermore, immunocompetent neonatal mice represent an interesting model, because key processes of brain development in rodent occur postnatally (while in humans they occur during the third trimester of fetal development)^27^. Besides, with the subset of animals that survive, sequelae associated with ZIKV infection may be further examined^15^. Taking these information into account, we infected 3-day old Swiss mice with ZIKV (2 x 10^7^ PFU). Indeed, this is a very high viral dose challenge, to the best of our knowledge the highest titers described to infected WT neonatal mice^22^. Treatments were carried out with sofosbuvir at 20 mg/kg/day, a dose administrated to mice during the preclinical development of this substance. The studies using this dosage supported, later on, the clinical dossier to approve the safe and effective use of sofosbuvir in humans at 400 mg/day to treat HCV infection^23^.

The virus challenge used was expected to cause significant mortality in untreated mice (indeed, less than 5 % of all the animals assayed survived). Therefore, we initially hypothesized that the expected outcomes with sofosbuvir would be an increase in the time of survival and/or reduction in mortality of infected animals. In fact, both expectations were achieved. Our results show that sofosbuvir treatment was associated with reduced mortality in infected mice, especially compared with animals that received no intervention. We identified an associated likelihood of lower mortality when comparing pre-treatment with late initiation of treatment (2 days after infection). This narrow and early time frame for antiviral intervention is common to other acute viral infections^28^. Mortality to pandemic influenza for example is reduced if neuraminidase inhibitors are administered early in the time course of infection, preferentially within 2.5 days after infection^28^. Sofosbuvir-treated animals not only had reduced mortality risk, but they also lived longer, even with our high dose viral challenge. Moreover, sofosbuvir prevented ZIKV mortality, suggesting its potential use for pre-exposure prophylaxis. This could be a desirable feature to treat pregnant women under in high risk areas of ZIKV infection.

ZIKV detection in kidney, urine, plasma, spleen before reaching out the brain^29-31^ has been associated as was a hallmark of the natural history of ZIKV infection. Sofosbuvir effects on survival were associated with an inhibition of acute virus infection and spread through different anatomical compartments. Since sofosbuvir reduced acute virus loads, sofosbuvir-treated ZIKV-infected animals displayed less foci of brain microhemorrhage; when compared to untreated counterparts. Moreover, sofosbuvir-treated mice responded properly to neuromotor reflex (righting); whereas untreated ZIKV-infected animals had severe impairment of this parameter.

On the long-term analysis of animals that survived, we noticed that ZIKV-infected animals had behavioral sequelae compatible with memory impairment. This is consistent with virus-induced cell death and cerebral inflammation in memory-forming areas^14,19,22,32,33^. Sofosbuvir-treated ZIKV-infected mice survived longer and in more numbers, than untreated animals. The surviving animals were also healthier– responding to memory testing behavior consistently with uninfected control mice.

Altogether, our results point out that sofosbuvir treatment at a pharmacological relevant concentration inhibits ZIKV replication *in vivo*, reducing mortality and blocking behavior sequelae in the short- and long-term analysis. These results are an important prove of concept and consistent with a prospective necessity of antiviral drugs to treat ZIKV-infected individuals.

## ACKNOWLEDGMENTS

Thanks are due to Drs. Carlos M. Morel, Marcio L. Rodrigues, Renata Curi and Fabrícia Pimenta for their strong advisement and support regarding the technological development underlying this project.

## AUTHOR CONTRIBUTIONS

Experimental execution and analysis - ACF, CZV, PAR, GB-L, YRV, MM, PPS and CS Provided critical material - KB

Data analysis, manuscript preparation and revision - ACF, CZV, PAR, GB-L, YRV, MM, PPS, CS, HCCFN, LC, AT, FAB, PTB and TMLS

Conceptualized the study – ACF, CZV, PAR, KB, FAB, PTB and TMLS All authors revised and approved the manuscript.

## Supplementary Video

Animals numbers

#1 – Sofosbuvir-treated ZIKV-infected mouse

#3 – Untreated ZIKV-infected mouse

